# Fine-scale fronts as hotspots of fish aggregation in the open ocean

**DOI:** 10.1101/2019.12.16.877571

**Authors:** Alberto Baudena, Enrico Ser-Giacomi, Donatella d’Onofrio, Xavier Capet, Cedric Cotté, Yves Cherel, Francesco d’Ovidio

## Abstract

Oceanic Lagrangian Coherent Structures have been shown to deeply influence the distribution of primary producers and, at the other extreme of the trophic chain, top predators. However, the relationship between these structures and intermediate trophic levels is much more obscure. In this paper we address this knowledge gap by comparing acoustic measurements of mesopelagic fish concentrations to satellite-derived fine-scale Lagrangian Coherent Structures in the open ocean. The results demonstrate unambiguously that higher fish concentrations are significantly associated with stronger Lagrangian Coherent Structures, and we observe that these features represent a limiting condition for high fish concentrations. A model, specifically built for mid trophic levels with realistic parameters, provides a possible mechanism of fish aggregation, and is coherent with the observations. These results may help to integrate intermediate trophic levels in trophic models, which can ultimately support management and conservation policies.

## Introduction

Marine biomass distribution is highly patchy and variable in time across the entire trophic web Bertrand et al. (2014); McManus and Woodson (2012). Discerning the factors underpinning ocean patchiness is fundamental to understand how they influence biogeochemical reactions and ecosystem stability Martin (2003); Lévy and Martin (2013). These issues are pivotal for conservation purposes Gaines et al. (2010) and for assessing the impact of climate change on the marine environment Hoegh-Guldberg and Bruno (2010).

One of the origins of the heterogeneity of the biotic fields is the dynamic nature of the ocean environments, which transforms the water masses on a large range of temporal scales, including those of ecological relevance. In this regard, a special role is fulfilled by the mesoscale and submesoscale McGillicuddy (2016), now commonly referred to together as “fine-scales”, which span a spatial range from a few to hundreds of kilometers.

One fruitful approach for capturing the structuring effect of fine-scale dynamics is the extraction of so-called Lagrangian Coherent Structures, and in particular Lagrangian fronts Haller (2015); Lehahn et al. (2018). Lagrangian Coherent Structures (LCSs) provide several types of information regarding flow properties, such as the location of fronts, barriers to transport Boffetta et al. (2001), or retentive and coherent regions d’Ovidio et al. (2013). One of the most common Lagrangian diagnostics used to determine LCSs is the Finite-size Lyapunov Exponent (FSLE, d’Ovidio et al., 2004). This measures the exponential rate of water parcel deformation and has maximal values over frontal regions.

By shaping and elongating water patches, Lagrangian Coherent Structures have been demonstrated to set the frontiers of phytoplanktonic patches in terms of chlorophyll concentration Lehahn et al. (2018), and even functional type d’Ovidio et al. (2010). This in turn enhances contacts between different communities, regulating plankton diversity De Monte et al. (2013).

More recently, advances in biologging programs provided evidence on the impact of fine-scale structures on top predators behavior. The concentration of predators foraging efforts has been observed in the neighbourhoods of Lagrangian fronts Kai et al. (2009); Scales et al. (2018). Furthermore, fronts detected by Lagrangian Coherent Structures (which in the following we will call Lagrangian fronts) have been observed to influence predators movements Della Penna et al. (2015). This could enhance energy transfer and gain Abrahms et al. (2018).

However, while the influence of Lagrangian fronts has been observed on both extremes of the trophic chain, much less is known about mid-trophic levels. Prants et al. (see in particular Prants et al., 2014) demonstrated a correlation between Pacific saury catches and Lyapunov exponents, and Watson et al. (2018) found that several fishery vessels track LCSs when targeting fishery spots. However, these results leave some concerns about possible biases because commercial fisheries provides only punctual observations, and because they are even known to use satellite images. Therefore, fishing vessels may target intentionally frontal systems. Unbiased fish measurements have been instead recently used by Sato et al. (2018) to analyse the relationship between a frontal system and acoustic measurements in a coastal upwelling system. This allowed the authors to highlight the different role played by in-shore and off-shore waters. In terms of the mechanisms which can explain how fine-scale structures influence mid-trophic biomass distribution, even less is known. Classical explanations are based primarily on bottom-up mechanisms along fronts with intense upwelling Yoder et al. (1994); Woodson and Litvin (2015). However, these hypothesis do not take into account the necessity of a maturation time, which in the case of fish is consistently longer than both the growth response of lower trophic levels and the lifetime of the front. Neither the fish behavior is considered, despite the fact that fish possess efficient sensorial and swimming capacities along the horizontal (Kasumyan (2004)and Supporting Information SI.2).

The objective of the present study is to analyze the relationship between fine-scale structures in the open ocean and mid-trophic organisms with unbiased, direct observations of fish concentrations, and to propose a mechanism by which fine-scales organize mid-trophic biomass.

This study was conducted in the subantarctic area of the southern Indian ocean. The functioning of this region is mainly regulated by the Kerguelen plateau, a major topographic barrier for the Antarctic Circumpolar Current (ACC). The plateau enriches in iron, a limiting nutrient, the high-nutrients-low-chlorophyll waters advected by the ACC. Depending on seasonal light conditions and stratification of the water column, this provokes a large annual phytoplanktonic bloom, which supports a rich trophic chain. This is one of the reasons for which the Kerguelen archipelago and its surrounding waters are part of one of the ten largest marine protected areas in the world (http://www.mpatlas.org/).

In this region, the myctophids, also known as lantern fish, are one of the most abundant groups of mesopelagic fish. They are also present in other oceans world-wide and are thought to constitute one the largest portions of world fish biomass Irigoien et al. (2014). They also represent important prey items for numerous predators Cherel et al. (2010). Myctophids are reported to play a central role in the carbon export to deep sea depths, and are suspected to affect the climate Kwon et al. (2009). Constituting a potentially massive harvestable resource, they are threatened to be exploited in the near future St. John et al. (2016). A better understanding of the mechanisms regulating their biomass is thus urgently needed.

In this paper, we relate acoustic measurements of fish concentrations and satellite derived diagnostics. These diagnostics allow us to identify the intensity of several fine scale fronts. The aim is to explore the degree to which fish distribution is shaped by these fine-scale features, and in particular Lagrangian fronts. Our results indicate that stronger Lagrangian fronts aggregate larger quantities of fish, although not all these structures are aggregation sites. We then propose a simple aggregating mechanism specifically designed for fish, based on a cue pursuing dynamic along the horizontal dimension. Myctophids are supposed to be able to orientate and to actively swim to search for food (hypothesis discussed in SI.2). They follow a gradient of a passive tracer, which is considered a proxy for heterogeneously-distributed zooplankton. All the parameters values of this model are set using observational data (SI.5) and none of them has been optimized nor fitted. The predictions of such model are consistent with the observed concentrations. In addition, the model provides also theoretical estimations of the dominant spatio-temporal scales of aggregation.

## Results

### Relationships between acoustic fish concentration and satellite-derived diagnostics

We used ship acoustic measurements acquired along 6 transects of 2860 linear kilometers. For each point of the boat trajectory, we computed a value of Acoustic Fish Concentration (AFC). This is representative of the fish concentration in the water column. Alongside this, we calculate two satellite derived diagnostics: the Finite-size Lyapunov Exponent (FSLE) and the Sea Surface Temperature (SST) gradient. These two metrics are typically associated with front intensity (see Materials and Methods for further details). Other diagnostics are reported in SI.1. In-situ temperature measurement were available only on a part of the transects, and were therefore not considered in the analysis.

Fig. 1 depicts two illustrative examples of the boat trajectory on August 29^*th*^, 2014 and August 31^*st*^, 2013 respectively. They are superposed on a field of Finite-size Lyapunov exponents. The latter are associated with frontal features. Each dot of the boat trajectory is colored proportionally to the AFC in that point. On both the panels, AFC values indicate a qualitative agreement with the FSLE, increasing in correspondence of the frontal features identified by high values of the Lyapunov exponents.

**Fig. 1:**
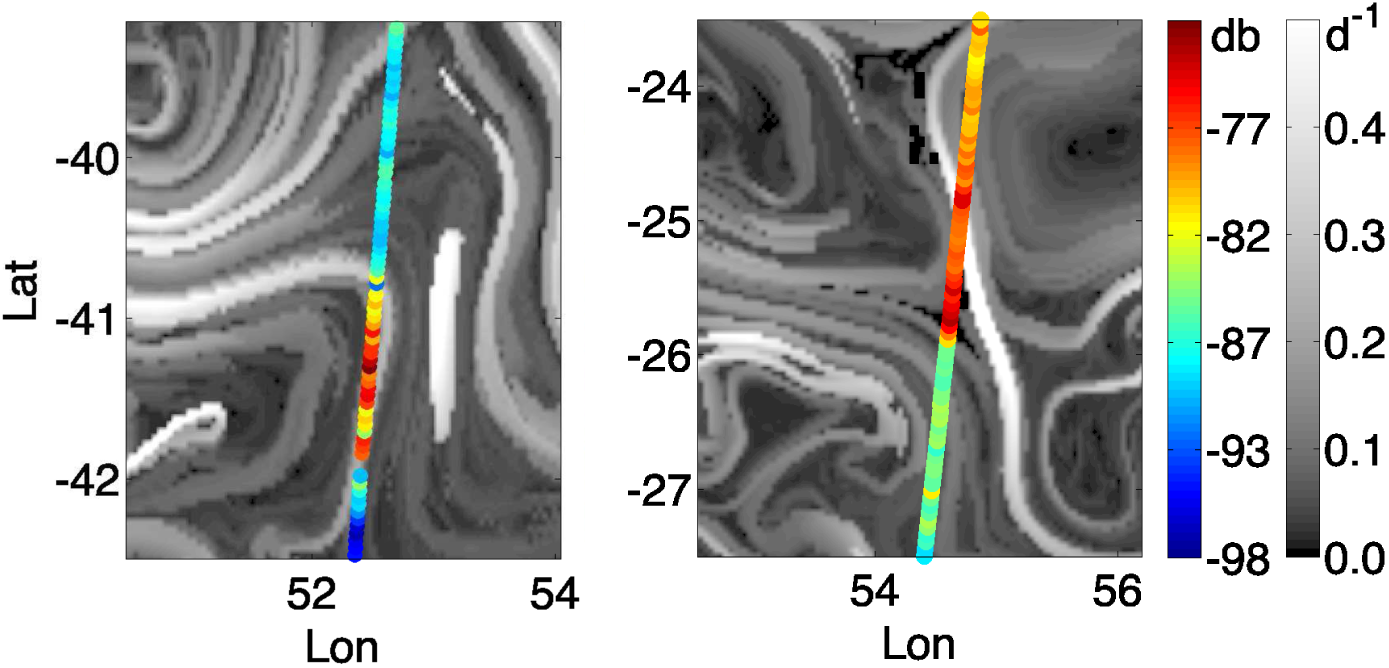
Illustrative examples of two transects of the boat trajectory. The color of each dot is proportional to the Acoustic Fish Concentration (in decibels, right colorbar for scale). The transects are superimposed on a simultaneous field of Finite-size Lyapunov Exponents. These identify fine-scale frontal structures. They are computed with altimetry derived velocities (days^−1^, left colorbar for the scale). Left panel: August, 29th, 2014. Right panel: August, 31st, 2013.

Fig. 2 depicts the scatter plots of the AFC against the two diagnostics, with both the axes in logarithmic scale. To determine whether the AFC values present significant differences in proximity of fine-scale features, a bootstrap analysis was conducted. Details of the methodology are provided in Materials and Methods section. The results are reported in Fig. 3. Significantly higher AFC values are detected over the thresholds (p-value < 0.001) for both FSLE and SST gradient. Finally, linear quantile regression method was employed Koenker (2005). This analysis was used to investigate the relationship between the higher values of AFC and the front intensity. Results of the regression are displayed in Fig. 2 as red lines. All the quantile slopes are statistically different from zero. This suggests the presence of a positive relationship between the higher values of AFC and FSLE and SST gradient (Table S.1).

**Fig. 2:**
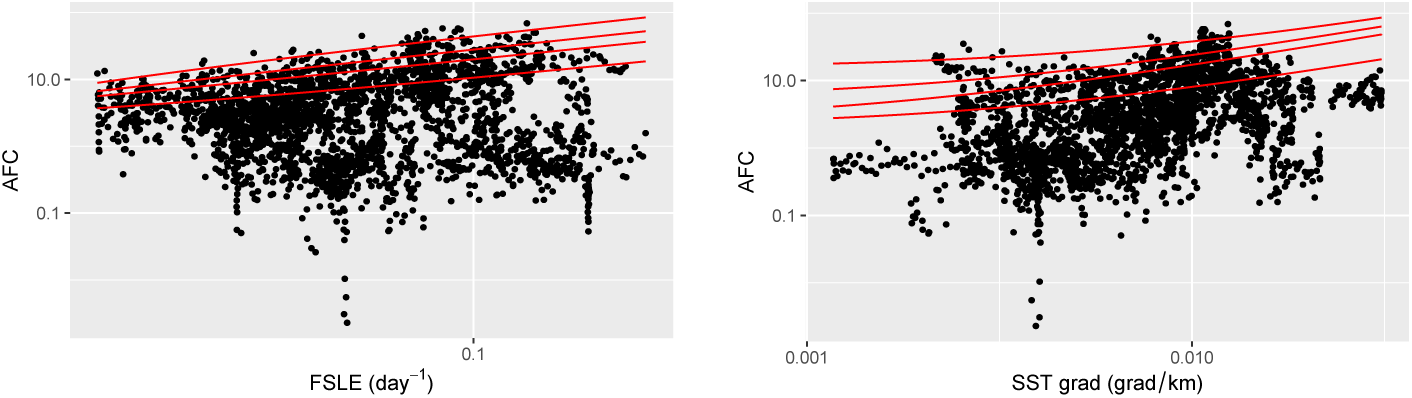
Scatter plot of AFC against FSLE (left panel) and SST gradient (right panel). The lines, from the bottom to the top, indicate the linear quantile regressions at 75th, 90th, 95th and 99th percentiles. The analysis is used to investigate the relationship between the front intensity and just the higher values of fish concentration. Both axes are in logarithmic scale (values equal to zero are therefore not depicted). Values of the quantile regression coefficients are reported in Table S.1.

**Fig. 3:**
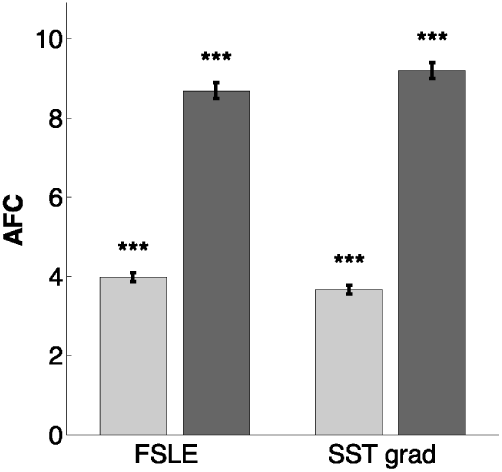
Bootstrap method results. Left columns refer to FSLE analysis, right columns to SST gradient. Light gray columns represent the mean AFC under the respective threshold, and the dark gray columns represent the mean AFC over the threshold. Error bars indicate the standard deviation, while black stars indicate the significance of the bootstrap test.

### A fine-scale mechanism of fish aggregation

Why do fish aggregate along frontal features? We addressed this question by proposing a simple mathematical model. The model assumes a gradient climbing capacity, which is one of the most widespread movement mechanisms used in other biological contexts (e.g., chemotaxis, Adler (1975)). This gradient climbing capacity is specifically tuned for mid-trophic organisms and myctophids and is based on a cue pursuing dynamic. Fish try to climb a gradient of tracer. We considered this tracer a proxy of zooplankton concentration, prey of several fish species, and in particular of myctophids Pakhomov et al. (1996). At the scales considered in this study (10s of kilometers) zooplankton swimming capacities are restricted to the vertical axis Genin et al. (2005). They can thus be considered, along the horizontal axis, passive tracers. Along this dimension, zooplankton aggregation and growth is usually driven by a relatively fast response to nutrients presence, of the order of days to weeks Vidal (1980). In particular, this is valid also for zooplankton species present in our study region Alonzo et al. (2003); Labat et al. (2005). Conversely, fish have growth rates consistently slower: in particular, pelagic fishes and myctophids are considered as “slow-growing fish”, with lifetimes spanning few years Greely et al. (1999). Therefore, their aggregation can not be explained by the same dynamics affecting the zooplankton. Alongside this, fish have extremely developed sensorial capacities and, differently from zooplankton, they can actively swim, with both capacities involved in many functional activities, including feeding Kasumyan (2004). These arguments support our approach of modelling zooplankton as a passive tracer and fish as active swimmers (we invite the reader to refer to SI.2 for further details). To include ocean patchiness, we perturbed the ability of the fish to properly identify the spots of zooplankton with a noise term. However, we assumed that fish are able to orientate without problems over a given threshold of the zooplankton gradient. This threshold was estimated from the zooplankton concentrations (SI.5).

Making these assumptions, the average velocity of a group of fish *U*_*F*_ (*x*) can be found applying simple algebra to a standard gradient climbing model (see Materials and Methods, subsection “Gradient climbing model”):

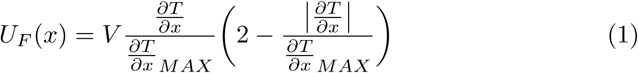

in which *T* is the zooplankton concentration, 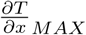 is the threshold of the zooplankton gradient, and V is the cruising speed of the myctophids.

Similarly, the evolution of the fish concentration over time can be easily obtained from the one-dimensional continuity equation, by imposing the conservation of fish total biomass, and by using Expr. 4 (see Materials and Methods for further details). The tracer shape used describes a sigmoid function, which models a generic local gradient in the zooplankton concentration. At the limits of the plateau, the tracer decreases along a distance of ∼5 km, a typical fine-scale range.

Four different snapshots (at 0, 6 hours, 1 day and 4 days) of the fish modelled concentration are illustrated in Fig. 4. Intuitively, having chosen a gradient-climbing behavior, one can expect that fish concentration will evolve quickly in the regions in which the tracer gradient is larger. Indeed, after only 6 hours, two peaks of doubled concentration are present in correspondence with the margins of the tracer plateau. In the following days, the two peaks decreased their growth rate, while the concentration between them increases until they merged together. The fish concentration is thus homogeneous over the plateau, presenting values between 2.5 and 3.5 times higher than the initial concentration. In case of a larger plateau (Fig. 5), the model predicts a similar behavior, but the merging between the peaks occurs over longer timescales. In that case, the peaks reach their maximum value after the first day, with the creation of two regions of increased concentration at the margins of the plateau. Changing the type of fish behavior leads to similar results (see SI.3).

**Fig. 4:**
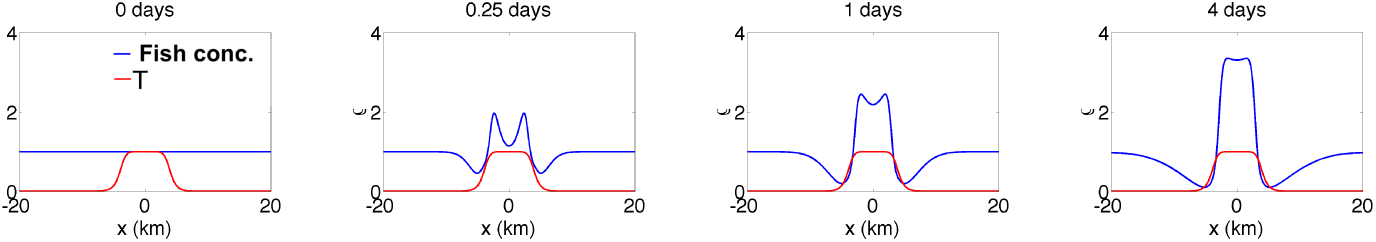
Time evolution of the fish concentration (blue line, adimensional) according to the continuity equation. The tracer (red line, adimensional) describes a plateau of 8 km in width. At its limits, its values range from 1 to 0 in about 5 km. Each panel represents a different snapshot at 0, 6 hours, 1 day, and 4 days.

**Fig. 5:**
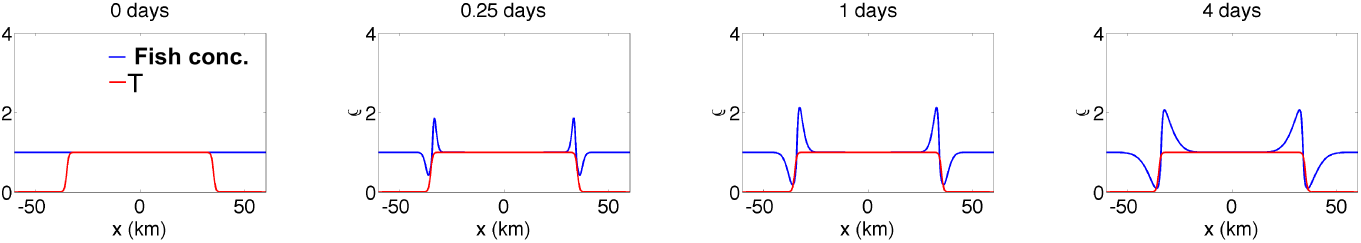
Time evolution of the fish concentration as reported in Fig. 4. This time, the plateau width has been set to 70 km.

We note that this mechanism does not explicitly take into account fronts. However, in the oceanic environment, frontal regions represent areas of convergence of water masses with different properties. This is why regions of strong gradients are often associated to frontal features Yoder et al. (1994).

However, at the same time during which this mechanism occurs, the tracer can evolve. As a sensitivity test, we numerically analyzed a scenario in which the tracer, subjected to a typical frontal dynamics, is stretched in a filament and eroded by diffusion (SI.4). Using realistic bio-physical parameters representative of the study area, we found that these tracer dynamics do not compromise the aggregation mechanism presented above, but may even facilitate the aggregation of fish along frontal zones. Indeed, the model developed predicts several quantitative information, such as an estimate of the fish aggregation over time. The latter highlighted a final concentration, on average, an order of magnitude stronger than the initial one. In addition, it was possible to obtain the “school life time”. This quantity indicates the amount of time during which a group of fish is able to follow a patch of interest before it vanishes due to the frontal dynamics. The school life times we obtained ranged between 7 to 25 days, with an average value of around 2 weeks. Remarkably, this amount of time matches with that of fine-scale processes.

## Discussion

In the oceans, patchiness characterizes biomass distribution, which regulates ecosystem stability and biogeochemical processes Bertrand et al. (2014); Martin (2003); Lévy and Martin (2013). In the open ocean, biomass distribution is largely driven by processes at the fine-scale (between 1 and hundreds of kilometers or a few days to several weeks, such as lateral advection and horizontal stirring Martin (2003); Lévy et al. (2018).

One approach to studying the structuring processing occurring at this scale is provided by Lagrangian methods. In the past, Lagrangian tools like the computation of Lagrangian Coherent Structures have revealed the structuring role of fine-scale dynamics on primary levels of the trophic chain (see a review in Lehahn et al. (2018)). On the other hand, they have highlighted their influence on apex predators behaviors as well Kai et al. (2009); Scales et al. (2018).

However, less information is available on intermediate trophic levels. In the present study (i) we correlate the intensity of acoustic fish concentrations and fine-scale fronts, and (ii) we propose a mechanism of fish aggregation along fronts. We analyze the Kerguelen region in the Southern Ocean, one of the ten largest protected areas in the world, and an oasis for several threatened species (IUCN, https://www.iucn.org/). The reference fish of this study are the myctophids, which are highly diffused in that region and one of the most abundant group of mesopelagic fish in the world Irigoien et al. (2014).

Our results reveal a significant difference in fish concentration between frontal and non-frontal features. These are difficult to explain with traditional mechanisms usually prescribed to lower trophic levels, such as rapid growth associated to a presence of nutrient, because fish have slower growth rates. Therefore, starting from a homogeneous fish distribution and applying simple ecological rules, we propose a gradient-climbing model specifically calibrated for the study of mid-trophic organisms. Once parameterized with values typical for the study region, the model demonstrates a quick response to gradient structures. Two peaks of doubled concentration appear after just 6 hours, and then merge after 4 days. Fish thus tend to be homogeneously distributed over the entire patch of interest. This is in coherence with on-going studies on myctophids’ response to food concentration. These presuppose that myctophids have a Holling type III functional response, in that they ingest always the same quantity of food over a certain prey density (A. Hulley, personal communication).

Tracer patchiness is known to be associated with frontal features Yoder et al. (1994); Lehahn et al. (2018). However, patchiness evolves, and under the double effect of stretching and diffusion, local gradients can be eroded. Thus, we tested the robustness of our model to this feature in SI.4. As in the former case, none of the parameters employed has been optimized nor fitted, but they all represent rigorous estimations of Southern Ocean physical and biological conditions, such as stretching and mixing rates or fish cruising speed. Results allowed us to estimate a typical lifetime for a fine-scale patch of around two weeks, much longer than the peak doubling time (∼ 6 hours). This robustness analysis is consistent with the hypothesis of a fixed tracer assumed in the gradient climbing scenario. Furthermore, we demonstrated that the stretching and diffusion dynamics can potentially enhance fish aggregations.

The patch of interest is considered as a proxy of zooplankton concentration. However, being parameterized as a passive tracer, it can be considered, more simply, as a physical water property. Indeed, physical characteristics, such as temperature, have been proven to be used by predators to find favorable conditions where concentrated food occurs Snyder et al. (2017).

The proposed mechanism of aggregation needs two obvious initial conditions: the presence of fish, and the presence of a zooplankton patch. It is presumable that fine-scale fronts, lacking one or both these conditions, cannot act as aggregating spots. Furthermore, while we assumed that the aggregation occurs after a certain amount of time, the environment studied is dynamic. Thus, it is likely that the aggregation mechanism was observed during different stages. The statistically significant positive relationships between the high values of acoustic fish concentration with the front intensity diagnostics confirms the previous considerations. Not all of the strong fronts detected indicate high acoustic fish concentrations. On the contrary, strong fish aggregation is preconditioned by the intensity of a frontal feature. Our results suggest that fronts represent in this regard a limiting condition for high fish concentrations. Model predictions are in accordance with the observations. Furthermore, the model provided estimations of other aggregation dynamics, such as the dimensions of the aggregation patterns or its intensity, with timescales comparable with those of fine scale processes. Specific experiments are necessary to validate these outputs, which however look promising.

Within the ACC, little information available on mid-trophic levels reported that the large circumpolar fronts are known to host (i) large densities of zooplankton and myctophids and (ii) that these organisms are patchily distributed Pakhomov and Froneman (2000). We identified in this study, at least partly, the potential mechanisms driving the patchiness observed at fine-scale. Assessing the preconditions and the other dynamics necessary for front aggregation is a new, open challenge emerging from the present work.

Note that in our work we focus only on open ocean Lagrangian fronts, that is, fine-scale frontal features induced by the mesoscale open ocean activity. In particular, we intentionally exclude coastal fronts as well as large scale fronts, whose dynamics and ecological role may be different (e.g., Lara-Lopez et al. (2012); Netburn and Koslow (2018); Sato et al. (2018)). Finally, the limitations of our analysis must be discussed. No vertical dynamics has been included in the model presented. However, these play an important role in the organization of marine biota Lévy et al. (2018), and, typically, stronger gradients are present along the vertical Ohman (1988). This assumption is due to the fact that, while knowledge on horizontal dynamics is more advanced, 3D Lagrangian analyses, while appealing Sulman et al. (2013), are not currently possible, due to a lack of quantitative information on the vertical velocities of the ocean. Future satellite missions (such as SWOT: https:/swot.cnes.fr) will possibly help to mitigate this problem, by helping the assimilation scheme to better reconstruct the three dimensional dynamics Morrow et al. (2019). At the same time, they will also improve satellite resolution, providing a more precise location of fine-scale fronts. The model studied does not consider the diel vertical migration of myctophids either. This choice is driven by the difficulty in parameterizing such a non linear behavior and in assimilating different migratory diel patterns. However, we note that many zooplankton species exhibit a diel cycle as well. Finally, zooplankton consumption due to fish foraging is considered to be negligible, or compensated by source terms (like blooms). The limitations presented can open the way for future investigations. Thus, the present study sheds some light on a largely unexplored topic.

The results presented here may be useful for improving the representation of intermediate trophic levels to coupled ecological and physical models Robinson et al. (2011), habitat models PC et al. (2018), targeting the mesopelagic compartment in particular. At the same time, the possibility of using Lagrangian Coherent Structures as a proxy of higher fish concentrations may further improve the integration of satellite-derived Lagrangian tools in conservation planning Penna et al. (2017).

## Materials and Methods

### Acoustic measurements

Two subsets of data were used for the analysis. Both of them were collected within the Mycto-3D-MAP program. The first subset (named MYCTO) was collected during 6 campaigns in 2013 and 2014, both in summer and in winter (see Tab. 1 for more details), using split-beam echo sounders at 38 and 120kHz. The data were then treated with a bi-frequency algorithm, applied to the 38 and 120 kHz frequencies (details of data collection and processing are provided in Béhagle et al. (2017)). This provides a quantitative estimation of the concentration of gas-bearing organisms, mostly attributed to fish containing a gas-filled swimbladder in the water column Kloser et al. (2009). Most mesopelagic fish present swimbladders and several works pointed out that myctophids are the dominant mesopelagic fish in the region Duhamel et al. (2014). Therefore, we considered the acoustic signal as mainly representative of myctophids concentration. Data were organized in acoustic units, averaging acoustic data over 1.1 km along the boat trajectory on average. Myctophid school length is in the order of tens of meters Saunders et al. (2013). For this reason, acoustic units were considered as not autocorrelated. Every acoustic unit is composed of 30 layers, ranging from from 10 to 300 meters (30 layers in total).

The second subset of data (named ZOO) was collected between January the 22^*nd*^ and February the 5^*th*^, 2014. It was processed in the same manner as the other dataset, but this time the bifrequency algorithm was used to infer the zooplankton biomass. This second subset possesses a higher resolution with an acoustic unit every 206 m of the boat trajectory on average.

The two datasets were used to infer the Acoustic Fish Concentration (AFC) and the Acoustic Zooplankton Concentration (AZC) in the water column, respectively. We considered as AFC (or AZC) of the point (*x*_*i*_, *y*_*i*_) the average of the bifrequency acoustic backscattering on the whole column, with the exclusion of the first layer. The latter was not considered due to surface noise.

The AZC was used to compute the zooplankton gradient. The zooplankton gradient of a point *i* of the boat transect is computed as:

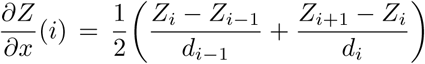

in which *d*_*i*_ indicates the kilometric distance between the point *i* + 1 and *i*, and *Z*_*i*_ is the cubic spline interpolation (to smooth the noise effects), in an around of 2 km, of the zooplankton concentrations. This type of interpolation, in contrast with the moving average, preserves the trend of the data, and thus a possible front.

**Table 1:** Details of the acoustic transects analyzed.

| Cruise | Season | St. Date | End Date | Distance (km) |
| --- | --- | --- | --- | --- |
| LOGIPEV193.RUNKER | Summer | 09/02/2013 | 17/02/2013 | 2752 |
| LOGIPEV193.KERMAU | Summer | 04/03/2013 | 10/03/2013 | 3781 |
| OP2013-2.RUNKER | Winter | 30/08/2013 | 10/09/2013 | 3310 |
| LOGIPEV197.RUNKER | Summer | 06/01/2014 | 13/01/2014 | 2800 |
| LOGIPEV197.KERMAU | Summer | 06/02/2014 | 18/02/2014 | 2045 |
| OP2014-2.RUNKER | Winter | 24/08/2014 | 04/09/2014 | 3677 |

### Regional data selection

The geographic area of interest of the present study is the Southern Ocean. To select the boat trajectory points belonging to this region, we used the ecopartition of Sutton et al. Sutton et al. (2017). Only points falling in the *Antarctic Southern Ocean* region were considered. We highlight that this choice is consistent with the ecopartition of Koubbi et al. Koubbi et al. (2011) (group 5), which is specifically designed for the myctophids, the reference fish of this study. Furthermore, this choice allowed us to exclude major fronts which have been the subject of different research works Lara-Lopez et al. (2012); Netburn and Koslow (2018). In this way our analysis is focused specifically on fine scale fronts.

### Day-night data separation

Several species of myctophids present a diel vertical migration. They live at great depths during the day (between 500 and 1000 m), ascending at dawn in the upper euphotic layer, where they spend the night. Since the maximal depth reached by our equipment is 300 m, in the analysis reported in Fig. 2 and 3 we excluded data collected during the day. However, their analysis is reported in SI.1. This is also consistent with the use of the Lagrangian and Eulerian diagnostics. These quantities, obtained from geostrophic velocity fields, are representative of the first part of the water column (∼ 50 m).

### Satellite data

#### Velocity current data and processing

Velocity currents are obtained from the Sea Surface Height (SSH), which is measured by satellite, through the geostrophic approximation. Data, which were downloaded from E.U. Copernicus Marine Environment Monitoring Service (CMEMS, http://marine.copernicus.eu/), were obtained from DUACS (Data Unification and Altimeter Combination System) delayed-time multi-mission altimeter, and displaced over a regular grid with spatial resolution of 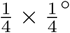 and a temporal resolution of 1 day.

Trajectories were computed with a Runge-Kutta scheme of the 4th order with an integration time of 3 hours. Finite-size Lyapunov Exponents (FSLE) were computed following the methods in d’Ovidio et al. (2004), with initial and final separation of 0.04° and 0.4° respectively. This Lagrangian diagnostic is commonly used to identify Lagrangian Coherent Structures. This method determines the location of barriers to transport, and it is usually associated with oceanic fronts Haller (2015).

#### Temperature field

The Sea Surface Temperature (SST) field was downloaded from the OSTIA global foundation Sea Surface Temperature product (SST_GLO_SST_L4_NRT_OBSERVATI The data are represented over a regular grid with spatial resolution of 0.05 × 0.05° and daily-mean maps.

#### Satellite data extrapolation

For each point of the boat trajectory, we considered as value of the diagnostic the average value over a disc of radius *σ*. Different *σ* were tested, and the best results were obtained with the *σ* = 0.2°, reference value reported in the present work. This value is consistent with the uncertainty of the satellite velocity field.

### Statistical processing

#### Bootstrap method

In this section, we describe the analysis necessary for the bootstrap test. To identify the frontal features, the following diagnostic values were chosen as thresholds for representing the front: for the FSLEs, we used 0.08 days^−1^, a threshold value consistent with those chosen in De Monte et al. (2012) and obtained from Kai et al. (2009). For the SST gradient, we considered representative of thermal front values greater than 0.9°C/100 km, which is about the half of the value chosen in De Monte et al. (2012). However, in that work, the SST gradient was obtained from the advection of the SST field with satellite-derived currents for the previous 3 days, which structures it in high resolution features that therefore present higher gradient values.

The threshold value splits the AFC into two groups: over and under the threshold. Their independency was tested using a Mann-Whitney or U test. Finally, bootstrap analysis is applied following the methodologies used in De Monte et al. (2012).

#### Linear quantile regression

Linear quantile regression method Koenker (2005) is employed to estimate the effect of FSLE and SST gradient on the upper limit of the AFC, as in Sankaran et al. (2005). The percentiles values used are 75th, 90th, 95th, and 99th. The analysis is performed using the statistical package QUANTREG in R (https://CRAN.R-project.org/package=quantreg, v.5.38, Koenker (2005)), using the default settings.

### Gradient climbing model

The fish searching dynamic is considered a one dimensional, individual-based, Markovian process. Time is discretized in timesteps of length Δ*τ*. We presuppose that, at each timestep, the fish chooses between swimming in one of the two directions. This decision depends on the comparison between the quantity of tracer at its actual position and the one perceived at a distance *f*_*W*_ from it, where *f*_*W*_ is the field view of the fish. We consider the fish immersed in a tracer whose concentration is described by the function *T* (*x*).

An expression for the average velocity of the fish, *U*_*F*_ (*x*), can now be derived. We assume that the fish is able to observe simultaneously the tracer to its right and its left. Fish sensorial capacities are discussed in SI.2. The tracer observed is affected by a noise. Noise distribution is considered uniform, with −*ξ*_*MAX*_ < *ξ* < *ξ*_*MAX*_, *ξ*_*MAX*_ > 0. The effective values perceived by the fish, at its left and its right, will be, respectively:

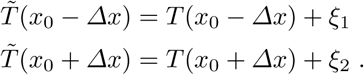

We assume that, if 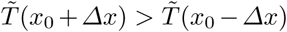, the fish will move to the right, and, vice versa, to the left. We hypothesize that the observational range is small enough to consider the tracer variation as linear, as illustrated in Fig. S.6 (SI.3). In this way:

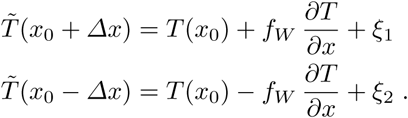

In case of absence of noise, or with 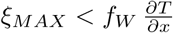, the fish will always move in the correct direction, in that it will climb the gradient. Assuming *V* as the cruising swimming velocity of the fish, this means *U*_*F*_ (*x*) = *V*. Let’s now assume 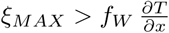. If *T* (*x*_0_ + Δ*x*) > *T* (*x*_0_ − Δ*x*) (as in Fig. S.6), and the fish will swim leftward if

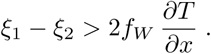

Since *ξ*_1_ and *ξ*_2_ range both between −*ξ*_*MAX*_ and *ξ*_*MAX*_, we can obtain the probability of leftward moving *P* (*L*). This will be the probability that the difference between *ξ*_1_ and *ξ*_2_ is greater than 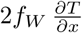

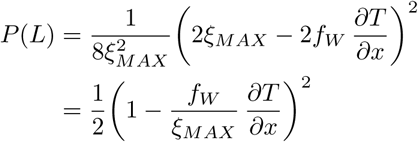

The probability of moving right will be

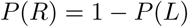

and their difference gives the frequency of rightward moving

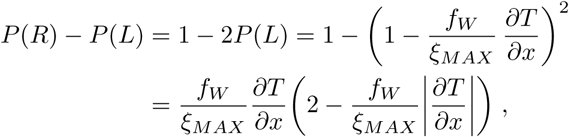

where the absolute value of 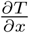 has been added to preserve the correct climbing direction in case of negative gradient. The above expression leads to:

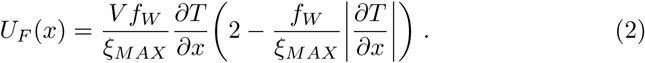

We then assume that, over a certain value of tracer gradient 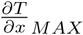,the fish are able to climb the gradient without being affected by the noise. This assumption, from a biological perspective, means that the fish does not live in a completely noisy environment, but that under favorable circumstances it is able to correctly identify the swimming direction. This means that

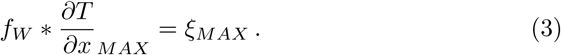

Substituting then (3) into (2) gives:

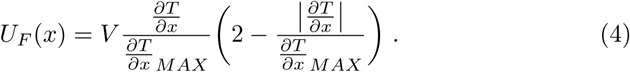

Finally, we can include an eventual effect of currents speed, considering that the tracer is transported passively by them, simply adding the current speed *U*_*C*_ to Expr. (4).

The representations provided are individual based, with each individual representing a single fish. However, we note that all the considerations done are also valid if we consider an individual representing a fish school. *U*_*F*_ will then simply represent the average velocity of the fish schools. This aspect should be stressed since many fish species live and feed in groups (see SI.2 for further details).

#### Continuity equation in one dimension

The solution of this model will now be discussed. The continuity equation is first considered in one dimension. The starting scenario is simply an initially homogeneous distribution of fish, that are moving in a one dimensional space with a velocity given by *U*_*F*_ (*x*).

We assume that in the time scales considered (few days to some weeks), the fish biomass is conserved, so for instance fishing mortality or growing rates are neglected. In that case, we can express the evolution of the concentration of the fish *ρ* by the continuity equation

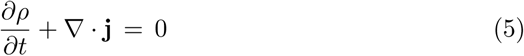

in which **j** = *ρ U*_*F*_ (*x*), so that Eq. (5) becomes

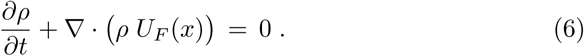

In one dimension, the divergence is simply the partial derivate along the *x-*axis. Eq. (6) becomes

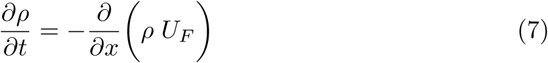

Now, we decompose the fish concentration *ρ* in two parts, a constant one and a variable one 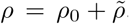. Eq. (7) will then become

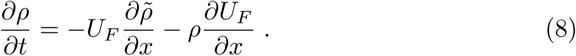

Using Expr. (4), Eq. (8) is numerically simulated with the Lax method. In Expr. (4) we impose that *U*_*F*_ (*x*) = *V* when *U*_*F*_ (*x*) > *V*. This biological assumption means that fish maximal velocity is limited by a physiological constraint rather than by gradient steepness. Details of the physical and biological parameters are provided in SI.5.

## Acknowledgements

This work is a contribution to the CNES/TOSCA project LAECOS and BIOS-WOT, and was partly funded by the Copernicus Marine Environment Monitoring Service (CMEMS) Sea Level Thematic Assembly Centre (SL-TAC). This work was supported financially and logistically by the Agence Nationale de la Recherche (ANR MyctO-3D-MAP, Programme Blanc SVSE 7 2011, Y. Cherel), the Institut Polaire Franais Paul Emile Victor, and the Terres Australes et Antarctiques Françaises. The authors would also like to thank Mara Baudena, Bettina Fach, Philippe Koubbi, Mark Ohman, Sara Sergi, Lars Stemmann and Jost von Hardenberg for their helpful advice.

## Authors contribution

A.B., E.S.G. and F.d.O designed the research with assistance from X.C. A.B. performed the research. A.B. and D.d.O. analyzed the data. C.C. and Y.C. provided the data. A.B. wrote the paper, with assistance from F.d.O., E.S.G, D.d.O, X.C., and C.C.

